# Accurate Macromolecular Complex Modeling for Cryo-EM with CryoZeta

**DOI:** 10.64898/2026.02.13.705846

**Authors:** Zicong Zhang, Shu Li, Farhanaz Farheen, Yuki Kagaya, Boyuan Liu, Nabil Ibtehaz, Genki Terashi, Tsukasa Nakamura, Han Zhu, Kafi Khan, Yuanyuan Zhang, Daisuke Kihara

**Affiliations:** Department of Computer Science, Purdue University, West Lafayette, Indiana, 47907, USA; Department of Biological Sciences, Purdue University, West Lafayette, Indiana, 47907, USA

**Keywords:** cryo-electron microscopy, cryo-EM, structural biology, protein structure modeling, protein structure prediction, diffusion model, multimodal deep learning

## Abstract

Cryogenic electron microscopy (cryo-EM) has become a widely used technique for determining the three-dimensional structures of biological macromolecules. Despite its advantages, building accurate structural models from cryo-EM data remains challenging, particularly at non-atomic resolutions. Here, we present CryoZeta, a de novo structure modeling program that leverages a diffusion-based generative deep neural network to integrate cryo-EM map density features with a biomolecular structure prediction pipeline similar to Alphafold3. By jointly leveraging sequence information and density-based features, CryoZeta generates highly accurate structural models that are consistent with the experimental map density. Evaluated on benchmark datasets covering protein complexes, protein–nucleic acid assemblies, and nucleic acid–only systems at resolutions up to 10 Å, CryoZeta consistently outperforms existing cryo-EM modeling methods in atomic accuracy. These results highlight the benefits of directly incorporating cryo-EM density into modern structure prediction pipelines and establish the method as a robust tool for automated, high-fidelity modeling from cryo-EM maps.

## Introduction

The three-dimensional (3D) structure of proteins and nucleic acids provides essential insights into the molecular mechanisms underlying their functions in living cells. Owing to several notable technical advantages^1,2^, cryogenic electron microscopy (cryo-EM) has recently become a primary method of choice for determining biomolecular structures.

Although an increasing number of cryo-EM maps are now determined at high resolution, interpreting them by building 3D structural models remains challenging when relying on conventional interactive tools^3–5^. When the resolution is worse than about 3 Å, complete main-chain tracing becomes difficult due to the presence of local regions with even lower resolutions. At about 5 Å or worse, identifying individual amino acids and nucleotides becomes problematic, and typically only portions of the structures can be modeled.

To facilitate structure modeling, various modeling tools have been developed^6–8^. Early protein modeling methods typically identify local dense regions in a map, which are likely to correspond to amino acid positions, and connect them to generate a main-chain trace^9,10^. Then, methods appeared that incorporate deep learning at the initial stage of modeling to identify amino acid positions in the map^11–14^. This use of deep learning has significantly enhanced modeling accuracy, as amino acid positions can now be detected more reliably. For nucleic acids, CryoREAD^15^ applies a procedure analogous to a protein modeling method, DeepMainmast^12^, in which nucleotide positions are first identified by a deep neural network and then connected to build RNA/DNA structures. These first-generation deep learning-based methods use deep learning for initial step for detecting atom positions, and then rely on conventional tracing approaches, such as traveling salesman solvers, other combinatorial optimization techniques, and dynamic programming for mapping the sequence, to build the main-chain trace.

More recently, modeling methods have been developed that use deep learning for both atom detection and the structure modeling step. ModelAngelo uses Graph Neural Network (GNN) to build a structure model, combining detected atom positions, cryo-EM features with a protein language model^16^. E3-CryoFold uses SE(3)-Equivariant GNN to combine local map features and a protein language model to build a structure model^17^. CryFold^18^, and SMARTFold^19^ use network architecture that is based on Alphafold2 (AF2)^20^. In their networks, predicted amino acid positions in a cryo-EM map are input into an attention module, where they are cross-referred with sequence-based embedding, and passed to a structure-building module to build full-atom protein models. As these methods are based on AF2, their applications are limited to proteins. Another limitation lies in the range of map resolutions these methods can handle. In most of these methods the neural networks were trained on maps with resolutions up to 4 Å, which partly explains why their structure modeling is effective only within that resolution range.

For maps of a worse resolution over about 5 Å, as de novo tracing is almost not possible, structure fitting methods have been developed as an alternative choice. Recent methods fit AF2 models to maps^21–23^. EMBuild^23^ operates on predicted residue positions derived from the map rather than the raw density map itself. DiffModeler^24^ uses a diffusion model^25^, a type of generative deep learning model, to enhance main-chain positions in cryo-EM maps, facilitating more accurate fitting of AF2 models.

Here, we present CryoZeta, the first deep learning framework that integrates spatial features from cryo-EM density maps into a diffusion model–based structure modeling pipeline that is similar to AlphaFold3^26^ (AF3). A key advantage of CryoZeta is that the diffusion-based approach inherently incorporates map density information during structure generation. This integration enables the seamless combination of experimental cryo-EM data with learned biomolecular sequence and structural features, resulting in self-consistent 3D models. Since CryoZeta builds on the AF3 framework, it is capable of modeling protein–nucleic acid complexes. Recently, CryoBoltz also employed the AF3 architecture for cryo-EM map–based structure modeling^27^; however, the work only shows preliminary results on three single-chain proteins with simulated maps for two of them, its broader applicability remains unclear.

We benchmarked CryoZeta on 200 cryo-EM maps with resolutions ranging from 1.98 Å to 10 Å, which consists of 169 protein-containing targets (targets with at least one protein chain), 55 nucleic acid (DNA/RNA) targets. CryoZeta achieved an average TM-score^28^ of 0.973 on protein-containing targets, outperforming existing methods including AF3 (TM-score: 0.871), DeepMainMast (TMscore: 0.847), and ModelAngelo (TMscore: 0.694). On the protein–nucleic acid complexes, CryoZeta also demonstrated strong performance with an average TM-score of 0.678 clearly exceeding AF3 (TM-score: 0.383) and ModelAngelo (TM-score: 0.174). The method is freely available as source code and through a web server at https://em.kiharalab.org.

### Overview of CryoZeta Architecture

CryoZeta takes protein and/or nucleic acid (DNA or RNA) sequences, together with a cryo-EM map, and builds a 3D structure consistent with the map density. We trained on maps with resolutions up to 8 Å and tested on maps with resolutions up to 10 Å. **Fig. 1** shows the overall deep neural network architecture of CryoZeta. It is primarily built upon the AF3 framework, which adopts a diffusion-based approach for structure generation. The architecture is inspired by SMARTFold^19^, which combined cryo-EM information into sequence-based structure prediction pipeline. While SMARTFold is based on AF2, limiting the application to proteins, CryoZeta is designed to handle proteins, DNA, and RNA, by adopting the AF3 architecture, enabling broader applicability to diverse biomolecular structures.

**Fig. 1.**
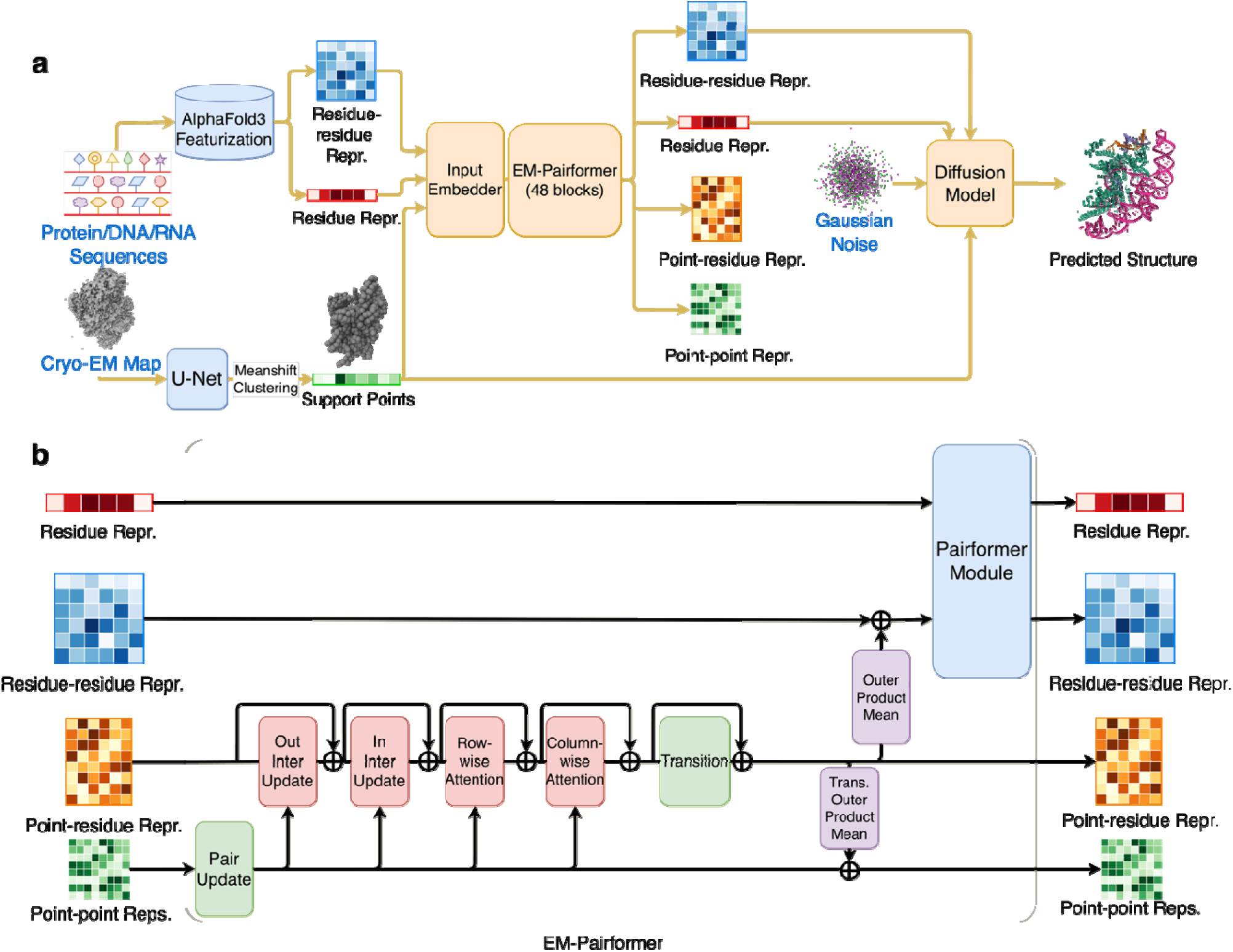
The neural network architecture of CryoZeta. **a.** The entire neural network pipeline. The network takes two types of input: protein or DNA/RNA sequences, and the cryo-EM map. The sequence is processed through the input data pipeline to construct an MSA from a database search, which is then used to produce residue and residue-pair representations. Meanwhile, atom positions are detected in the cryo-EM map using a 3D U-Net and subsequently clustered. The resulting positions, termed support points, are embedded into a point-pair representation using geometric features. These support points, together with the single- and pair-representations, serve as input to EM-Pairformer. EM-Pairformer operates on four representations: single, residue/nucleotide pair, support point–residue pair, and support point–support point. These representations interact and are jointly updated via cross-attention within EM-Pairformer. The updated single and residue/nucleotide pair representations are then passed to the diffusion-based structure module, which generates the 3D atomic structure through iterative denoising. In parallel, a point–residue/nucleotide distogram head predicts distances between residues (or nucleotides) and support points, providing spatial constraints that guide fitting of the predicted structure into the EM map. **b**. More detailed view of EM-Pairformer. The network within the transparent orange box corresponds to EM-Pairformer in panel a. In the complete architecture, 48 EM-Pairformer blocks are stacked.

The input to CryoZeta is protein/DNA/RNA sequences and cryo-EM density map. The input sequences are used to search for similar sequences from a sequence database, from which multiple sequence alignments (MSAs) are generated and represented as input embeddings. MSAs for query sequence were generated through the ColabFold MSA pipeline^29^, which collects sequences from UniRef100^30^ and ColabFoldDB, a non-redundant metagenome sequence database. For RNA, we employed rMSA pipeline^31^, which searches in RNAcentral^32^, Rfam^33^, and NCBI nucleotide (nt) database^34^. We did not generate MSAs for DNA sequences. Structure templates are not used in our framework. In the cryo-EM map, atoms of proteins and nucleic acids, namely, main-chain heavy atoms (N, Cα, C, O), Cβ, and other side-chain atoms from proteins and C1′, sugar, phosphate, and base atoms in nucleic acids are detected using a dedicated network composed of a hybrid convolution-attention based 3D U-Net model^35^ (**Supplementary Information 1**, **Supplementary Fig. S1, S2**). Based on the predicted atoms, amino acid and nucleotide types are also predicted using a second U-Net model (**Supplementary Fig. S1**). The identified Cα atoms of amino acids and C1′ atoms of nucleotides are clustered using the mean-shift algorithm^36^, and the resulting points are referred to as support points. As will be discussed below, we developed a variant of CryoZeta with support point interpolation where we generated five interpolation points between every pair of support points and assigned each point with the predicted probability of Cα/C1′ atom of the nearest grid point in the cryo-EM map. These support points, together with single- and pairwise embeddings, are input information that is processed by a module named Input Embedder (**Supplementary Information 2**) followed by another module named EM-Pairformer (**Fig. 1b**, **Supplementary Information 3**). The EM-Pairformer module takes four inputs: (i) target sequence embeddings, (ii) residue–residue representations that capture distance relationships between amino acid and nucleotide positions, (iii) support point representations derived from the cryo-EM map, and (iv) support point–residue representations that encode the correspondence between residues/nucleotides and support points in the map. These representations are iteratively updated through attention mechanisms to maintain consistency across them. The EM-Pairformer consists of 48 stacked blocks, each with independent parameters. The resulting residue and residue–residue representations are then passed to the diffusion model (**Supplementary Information 4**), which generates the 3D structure. The output structure from the diffusion model is recycled up to *x* times, in the form of residue–pair distances and residue representations, a procedure that typically improves the accuracy of the predicted structure. For more technical details of the network, see Supplementary Information 1 to 4. The inference time for a macromolecule with a total of 1,500 residues/nucleotides is less than 25 minutes (**Suppl. Fig. S3**).

Three networks were trained for CryoZeta: two 3D U-net networks that detect atoms and residue types (amino acids in proteins and nucleotides in DNAs and RNAs), respectively, in cryo-EM maps (**Fig. 1a**) and the end-to-end network that process input information, EM-Pairformer, and the diffusion model in **Fig. 1a**, which produces macromolecular structure models for the input cryo-EM maps. Training, validation, and testing the networks were performed on datasets collected from Protein Data Bank (PDB)^37^ entries determined by cryo-EM and thus have corresponding cryo-EM maps in Electron Microscopy Data Bank (EMDB)^38^.

The U-net for amino acids was trained on a training dataset of 5,917 maps (**Supplementary Table 1**) and validated on a set of 186 maps (**Supplementary Table 2**), where maps across the two datasets are non-redundant to each other. The maps underwent preprocessing, which included resampling the map voxels to 1Å with trilinear interpolation and normalization of their density values between 0 and 1. Then for each grid position of the map, labels were assigned based on the corresponding PDB files. For the atom detection network, we assigned labels of eight atom classes (Cα, backbone, Cβ, side chain atoms for proteins and C1′, sugar, phosphate, and sugar atoms for nucleic acids) to grid points. Structure modeling by CryoZeta model was validated on 186 maps (**Supplementary Table 3**) that are clustered into 65 groups with 40% sequence identity cutoff, following AF2^20^. Atom detection by U-nets and the structure modelling by the entire CryoZeta model was tested on 200 maps in 200 clusters (**Supplementary Table 4**). For more details, see Methods.

## Results

### Structure modeling performance of CryoZeta

First, we discuss the summary of structure modeling results from various angles (**Fig. 2**). Individual data is provided in **Supplementary Table 5-13.** The analysis in **Fig. 2** starts with two panels, that report the accuracy of atom (**Fig. 2a**) and residue/nucleotide detection (**Fig. 2b**) in cryo-EM maps. The hit rate plotted here is the fraction of atoms that were correctly detected within 2 Å among all the atoms in the reference structures in PDB. Cα atoms in proteins and C1′ atoms in nucleic acids, which we used for structure modeling, were detected with a high hit rate of 0.797, and 0.657, respectively, among other atoms. Compared to DeepMainmast^12^, CryoZeta detects Cα with a higher rate although direct comparison is not possible because different datasets were used. The trend that Cα is detected more reliably than Cβ is consistent across both studies. Backbone (BB) and sugar (SGR) atoms exhibited lower hit rates overall, but we leveraged their detected positions to improve Cα and C1′ atom detection for maps with resolutions worse than 5 Å, as described in **Supplementary Fig. S4 and S5**. These atom classes provide complementary information that is particularly beneficial at lower resolutions. In **Fig. 2b**, the average hit rate of amino acids and nucleotides were 0.528 ± 0.278 and 0.498 ± 0.266, respectively. Amino acids with high hit rates include those with aromatic rings, as well as proline and glycine, which have distinctive chemical structures. Detection performance based on voxels are provided in **Supplementary Fig. S6 and S7**.

**Figure 2.**
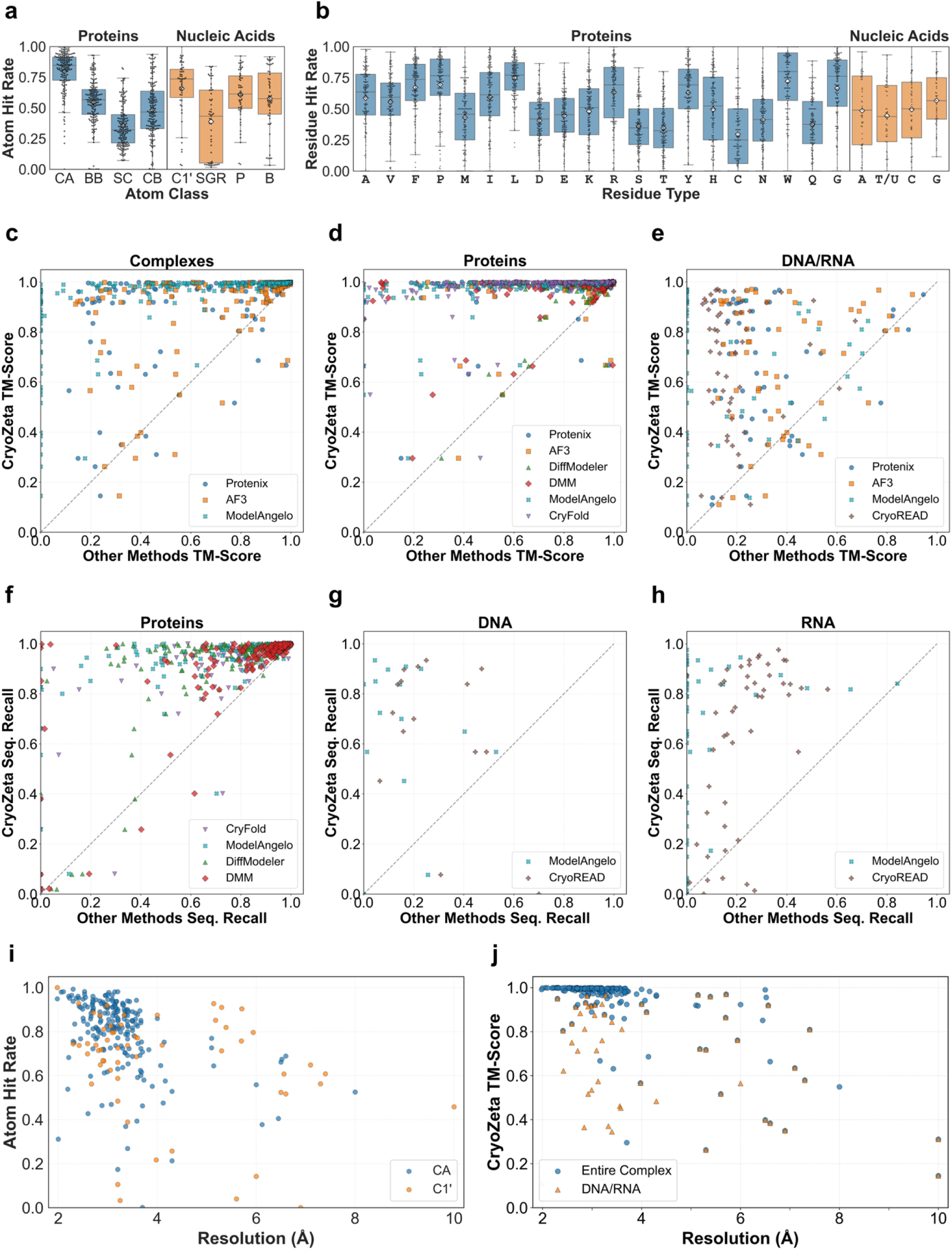
Performance of CryoZeta across atom-, residue-, and structure-level metrics for proteins, nucleic acids, and complexes. **a,** atom detection accuracy of four atoms each for proteins (Cα, backbone, side-chain, Cβ) and for DNA/RNA (C1′, SuGaR, Phosphate, Base). 200 test maps were plotted. Box plots were used to describe the distributions, with boxes spanning the first to third quartiles. Whiskers extend to 1.5 × the interquartile range. The median is shown as a line in the middle of the box, and the mean is indicated using a white diamond. The mean value of the eight atoms was Cα: 0.797; backbone: 0.547; side-chain: 0.357; Cβ: 0.485; C1′: 0.657; SGR: 0.389; Phosphate: 0.606; base: 0.576. For more detailed definition of the atom hit rate, see text. **b,** Amino acid and nucleotide detection accuracy. T in DNA and U in RNA were detected by the same class. The mean values were: 0.587, 0.554, 0.672, 0.695, 0.43, 0.593, 0.75, 0.407, 0.444, 0.48, 0.633, 0.361, 0.346, 0.63, 0.495, 0.296, 0.411, 0.721, 0.375, 0.665 for the 20 amino acids in the order shown in the plot (A, V, F, P, M, I, L, D, E, K, R, S, T, Y, H, C, N, W, Q, G) and 0.485, 0.444, 0.496, 0.567 for the 4 nucleotide classes (A, T/U, C, G), respectively. **c.** TM-score of protein/DNA/RNA complex models by CryoZeta in comparison with Protenix (blue circle), AF3 (orange square), and ModelAngelo (cyan cross). TM-score evaluates the global structure of a structure model. 200 targets in the test set were plotted. **d**, TM-score of protein models by CryoZeta in comparison with six methods, Protenix (blue circle), AF3 (orange square), DiffModeler (green upward triangle), DeepMainmast (DMM; red diamond), ModelAngelo (cyan cross), and CryFold (purple downward triangle). 169 protein models in the test set were plotted. If a target includes DNA/RNA, only the protein chains were evaluated. **e**, TM-score of DNA/RNA models by CryoZeta compared with Protenix, AF3, ModelAngelo, and CryoREAD (purple plus). 55 models in the test set were plotted. If a target includes proteins, only DNA/RNA chains were evaluated **f**, Sequence recall of protein models in comparison with four methods, CryFold, ModelAngelo, DiffModeler, and DMM. Sequence recall evaluates positions of amino acids and nucleotides in the model in the map. **g**, Sequence recall of DNA models in comparison with ModelAngelo and CryoREAD. 15 targets that have DNA in the test set were plotted. **h**, Sequence recall of RNA models. 48 targets that have RNA in the test set were plotted. **i,** Atom hit rate relative to the global map resolution. 207 maps with Cα atoms and 59 maps with C1′ atoms were plotted. **j,** TM-score of models relative to map resolution. 200 maps with a complex structure and 55 maps with DNA/RNA models were plotted.

The next three panels, **Fig. 2c–e**, show the global accuracy of structure models in terms of TM-score^39^. CryoZeta was compared with six existing methods, Protenix, AlphaFold3 (AF3), DiffModeler, DeepMainmast, ModelAngelo, and CryFold. Some methods were excluded from the complex structure evaluation (**Fig. 2c**) and the DNA/RNA structure evaluation (**Fig. 2e**) if they are not capable of modeling DNA/RNA structures. CryoREAD was included only in the DNA/RNA structure comparison (**Fig. 2e**), as it is specifically designed for those targets. For complex (**Fig. 2c**) and protein **(Fig. 2d**) structure evaluation, CryoZeta clearly outperformed the other methods. CryoZeta modeled 85.5 % (171 out of 200) of complex targets with a TM-score greater than 0.9. The average TM-score of CryoZeta was 0.933 for complexes and 0.973 for proteins, substantially higher than AF3, the method with the second-highest scores, which achieved 0.794 and 0.871, respectively. There are two targets in the protein evaluation (**Fig. 2d**) where CryoZeta’s model had a lower TM-Score than those built by AF3, Protenix, DeepMainMast, Diffmodeler. One of the targets is *Tribolium castaneum* ABCH-9C in complex with ceramide (EMD-39697, PDB: 8YZP). This protein forms a homodimer, and CryoZeta predicted a domain-swapped structure, resulting in a TM-score of 0.667, compared with other methods (AF3: 0.964, Protenix: 0.981, DeepMainmast: 0.995, DiffModeler: 0.968). Another target is the *Alkalihalobacillus halodurans* trp RNA-binding attenuation protein (Aha TRAP; EMD-44473, PDB: 9BE8), which assembles as a homohexamer forming a ring structure. CryoZeta split three of the chains into two connected domains, while correctly modeling the remaining three chains as intact units. This led to a TM-score of 0.686, compared with other methods (AF3: 0.987, Protenix: 0.869).

CryoZeta also maintained its superiority in DNA/RNA modeling (**Fig. 2e**). The average TM-score achieved by CryoZeta for DNA/RNA structures was 0.678, nearly double that of AF3 (0.383). ModelAngelo had a TM-score of 0 for 32 targets, producing highly fragmented models that are not meaningful.

The subsequent three panels (**Fig. 2f–h**) evaluate sequence recall, which measures model–map agreement at the amino acid and nucleotide levels. In these evaluations, Protenix and AF3 were not included because they do not provide model orientations relative to the map. Whole complex models were not evaluated, as the software used for this analysis, *phenix.chain_comparison*^40^, evaluates proteins and DNA/RNA separately. Consistent with the TM-score results, CryoZeta again outperformed the other methods in this metric. For protein targets, the second-best method was DeepMainmast, which achieved a sequence recall of 0.830, compared with 0.957 for CryoZeta. **Supplementary Fig. S8** shows the backbone recall for the same set of methods, which evaluates whether atoms in the reference structure are modeled within 3 Å, without considering atom identity or residue/nucleotide type. This analysis yields results consistent with the sequence recall results.

The last row (**Fig. 2i, j**) examines the atom hit rate and TM-score of CryoZeta as a function of global map resolution. The Cα detection rate gradually decreased with worsening map resolution (Fig. 2i). C1′ atom detection also declined at lower resolutions; however, due to the augmented C1′ detection procedure (**Supplementary Fig. S5**), the decrease was less pronounced than for Cα atoms. In contrast, TM-score (**Fig. 2j**) showed a weaker dependence on resolution. Notably, 94.0% of complexes derived from maps at 4 Å resolution or better achieved a TM-score greater than 0.9.

In **Table 1**, we summarize the modeling performance of CryoZeta in comparison with other methods. CryoZeta outperformed all existing methods across all three metrics and categories. Notably, CryoZeta maintained superior modeling performance even for low-resolution maps worse than 5 Å, achieving a TM-score of 0.694, compared with 0.548 for AF3, 0.448 for Protenix, and 0.003 for ModelAngelo.

**Table 1.** Performance comparison of cryo-EM structure modeling methods across complexes, proteins, and DNA/RNA targets. Global structural accuracy is reported using TM-score for complexes, proteins, and DNA/RNA. Backbone (BB) recall and sequence recall quantify model–map agreement at the residue and nucleotide levels for proteins and nucleic acids, respectively. Methods that do not support specific target types or do not provide model–map orientation were excluded from the corresponding evaluations and are indicated by dashes. The number of targets are as follows: complexes: 200; proteins: 169; DNA/RNA: 55; DNA: 15; RNA: 48. Evaluation values were assigned a score of 0 if a method failed to produce a structure due to low map resolution or excessive fragmentation, preventing the evaluation software from assessing agreement with the reference structure. CryoZeta variants (“Interpolate” and “Combine”) represent alternative atom-probability integration strategies evaluated on the same benchmark dataset. In each column, the highest value is highlighted in bold.

| Method | TM-score |  |  | BB Recall |  |  | Seq Recall |  |  |
| --- | --- | --- | --- | --- | --- | --- | --- | --- | --- |
|  | Complexes | Proteins | D/RNA | Proteins | DNA | RNA | Proteins | DNA | RNA |
| Protenix | 0.758 | 0.836 | 0.325 | - | - | - | - | - | - |
| AF3 | 0.794 | 0.871 | 0.383 | - | - | - | - | - | - |
| DiffModeler | - | 0.897 | - | 0.844 | - | - | 0.757 | - | - |
| DeepMainMast | - | 0.847 | - | 0.881 | - | - | 0.830 | - | - |
| CryFold | - | 0.689 | - | 0.853 | - | - | 0.828 | - | - |
| ModelAngelo | 0.578 | 0.694 | 0.174 | 0.774 | 0.78 | 0.26 | 0.761 | 0.15 | 0.08 |
| CryoREAD | - | - | 0.190 | - | 0.71 | 0.53 | - | 0.29 | 0.23 |
| CryoZeta | 0.933 | 0.973 | 0.678 | 0.957 | **0.79** | 0.72 | 0.929 | 0.66 | 0.63 |
| CryoZeta-Interpolate | 0.927 | 0.969 | 0.673 | **0.964** | 0.77 | 0.70 | **0.935** | 0.65 | 0.62 |
| CryoZeta-Combine | **0.937** | **0.976** | **0.695** | **0.964** | **0.79** | **0.75** | 0.934 | **0.69** | **0.65** |

For CryoZeta, results from two method variants are also reported. In CryoZeta-Interpolate, five additional interpolated points are introduced between each pair of support points to guide downstream structure modeling. This interpolation is motivated by the observation that a major source of modeling failure arises from insufficient atom detections in the map. CryoZeta-Combine selects the model with the highest model reliability score (Eq. 1) from five candidate models generated by CryoZeta and CryoZeta-Interpolate. The model reliability score is defined as follows:

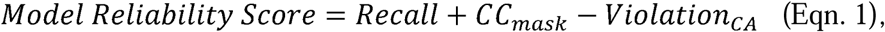

where Recall measures the fraction of predicted Cα/C1′ atoms that have at least one support point within 3Å, *CC_mask_* is the cross-correlation coefficient computed between the simulated density derived from the structure and the experimental cryo-EM map, with the calculation restricted to voxels within a defined region surrounding the molecular structure, *Violation_cA_* is defined as the number of Cα–Cα distances between consecutive residues that exceed 4 Å. The violation term is applied only to predicted protein structures, because CryoZeta occasionally produces domain-swapped conformations that exhibit large Cα–Cα distance violations.

CryoZeta achieved higher average performance than CryoZeta-Interpolate across all evaluated metrics. However, on individual targets, CryoZeta-Interpolate often produced higher TM-scores than CryoZeta, in some cases by a substantial margin (**Supplementary Fig. S9**). By leveraging the model reliability score (Eq. 1), CryoZeta-Combine is able to select the better model in most cases, resulting in consistent performance improvements across all metrics in Table 1.

### Modeling examples of protein complexes

In this section, we highlight five illustrative examples of CryoZeta’s structure models. We start with five examples of protein complex models (**Fig. 3**). The first example (**Fig. 3a**) is the SpoIVFB–pro-σK complex (EMD-43288; PDB 8VJL), a hetero-octameric assembly comprising 1,796 residues. CryoZeta reconstructed the complete complex with the highest modeling accuracy among the compared methods (TM-score: 0.995). In contrast, AlphaFold3 produced a markedly different and more compact assembly, resulting in a substantially lower TM-score (0.460). ModelAngelo generated a partially correct model (TM-score: 0.850) but failed to model four β-hairpin regions (residues 251–261 of stage IV sporulation protein FB; red arrows), which correspond to regions of locally low map density.

**Figure 3.**
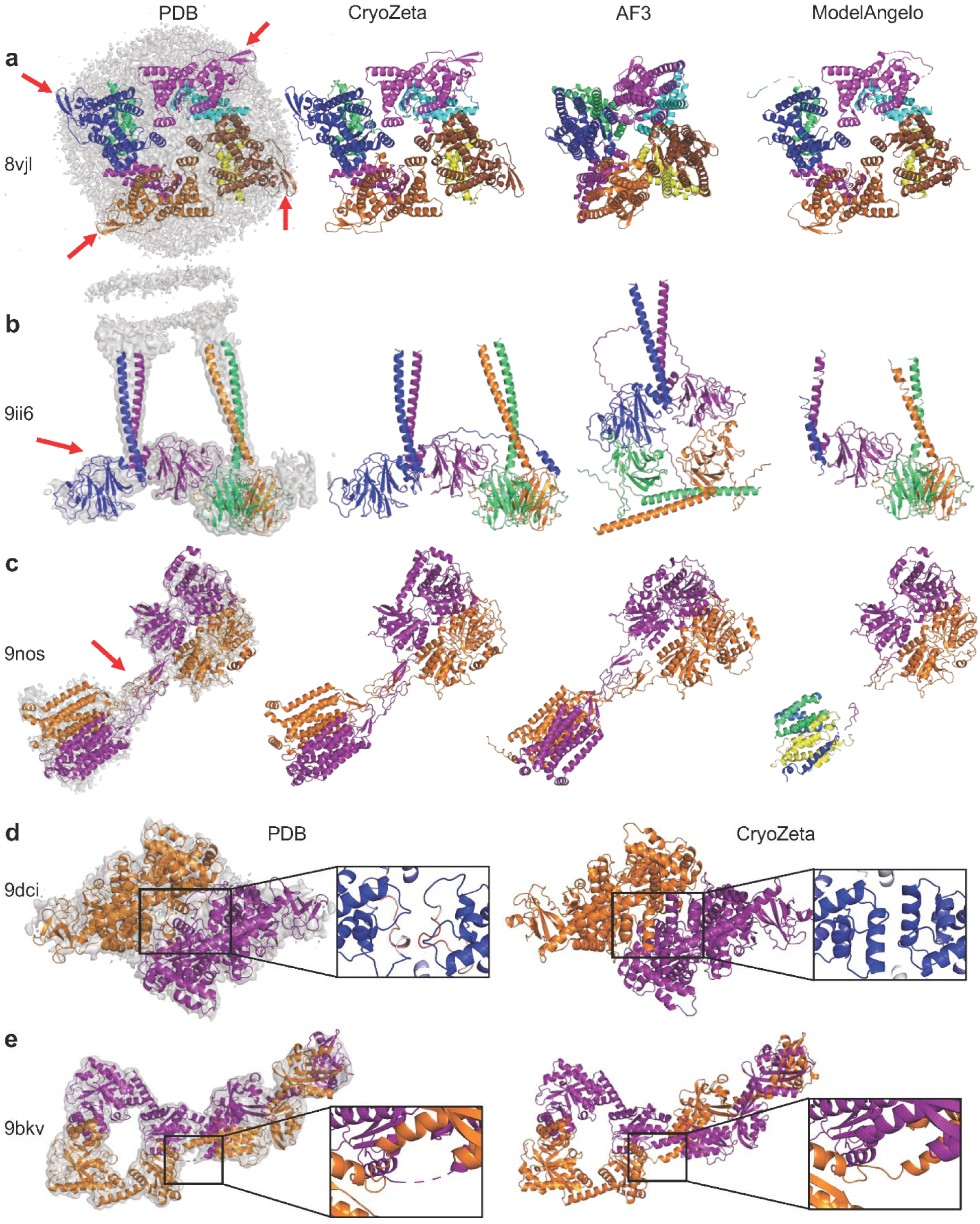
Examples of modeling for protein complexes. For each target, the cryo-EM density map at the recommended contour level, the reference PDB structure, and models predicted by different methods are shown. Modeling accuracy is assessed using TM-score, computed by comparison with the reference PDB structure. Different chains are displayed in distinct colors. **a.** SpoIVFB:pro-sigmaK complex (EMD-43288, res. 3.50□Å; PDB 8VJL), a hetero-octamer complex with 1,796 residues. TM-scores are CryoZeta: 0.995; AF3: 0.460; and ModelAngelo: 0.850. **b.** Intracellular domains of the BTN2A1–BTN3A1–BTN3A3 complex (EMD-60591, res. 3.27□Å; PDB 9II6), a hetero-tetramer complex with 1,057 residues. TM-scores are CryoZeta: 0.989; AF3: 0.433; and ModelAngelo: 0.728. **c.** Human sweet taste receptor (EMD-49612, res. 3.50□Å; PDB 9NOS), a hetero-dimer complex with 1,707 residues. TM-scores are CryoZeta: 0.997; AF3: 0.754; and ModelAngelo: 0.576. The ModelAngelo model is fragmented, with each fragment shown in a different color. **d.** Mycobacterium tuberculosis UvrD1 dimer (EMD-46752, res. 4.07□Å; PDB 9DCI), a homo-dimer with 1,542 residues. TM-score is CryoZeta: 0.941. Boxes show a magnified view of the dimer interface (residues 438-456), colored by DAQ score. Negative and positive DAQ scores are shown in red and blue, respectively. A higher DAQ score indicates stronger local density support for the assigned amino acid type. The reference structure (PDB 9DCI) lacks helical secondary structure in this region and shows negative DAQ scores. The CryoZeta model forms α-helices that have positive DAQ scores. **e.** DosP R97A bent-form dimer (EMD-44646, res. 3.43□Å; PDB 9BKV), a homo-dimer complex with 1,614 residues. The CryoZeta model achieves a TM-score of 0.631. The overall architecture of the predicted model closely matches the reference structure. However, a domain swap is observed at residues 384-388, highlighted in the boxes.

**Fig. 3b** shows the intracellular domains of the butyrophilin family BTN2A1–BTN3A1–BTN3A3 complex (EMD-60591; PDB 9II6), a hetero-tetramer comprising 1,057 residues. CryoZeta accurately reconstructed the overall complex structure with a TM-score of 0.989. Among the compared methods, only CryoZeta and AF3 correctly modeled the BTN3A1 subunit (blue chain, indicated by the red arrow); however, AF3 failed to assemble the complete complex. The next example is the human sweet taste receptor (EMD-49612; PDB 9NOS), a heterodimeric complex comprising 1,707 residues (**Fig. 3c**). CryoZeta achieved near-perfect agreement with the reference structure (TM-score: 0.997), including accurate modeling of the cysteine-rich (CR) domain, a flexible linker between the extracellular Venus flytrap agonist-binding (VFT) domain and the transmembrane (TM) domain (red arrow). This region was not correctly modeled by the other methods. In addition, the AF3 model exhibited an incorrect orientation of the seven transmembrane helices, resulting in a lower TM-score of 0.754. ModelAngelo was unable to model the cysteine-rich (CR) domain; moreover, the spatial arrangement of helices in the transmembrane and Venus flytrap (VFT) domains were inaccurate in several places.

For the *Mycobacterium tuberculosis* UvrD1 homodimer (EMD-46752; PDB 9DCI; **Fig. 3d**) at lower resolution (4.07 Å), CryoZeta achieved a TM-score of 0.941, with only minor structural differences relative to the reference PDB structure at the dimer interface (residues 438-456; boxed). CryoZeta modeled α-helical secondary structure in this region, whereas the deposited reference structure lacks helical features. Notably, DAQ validation scores^41^, visualized by the blue-to-red color scale in the boxed regions, indicate that the α-helices modeled by CryoZeta are better supported by the cryo-EM density (average DAQ score: +1.06, while the corresponding region in the PDB structure shows negative DAQ values (-0.09), suggesting insufficient density support. These results indicate that the CryoZeta model is more consistent with the experimental density and is likely to be more accurate in this region.

Finally, we present a challenging case for CryoZeta: the DosP R97A bent-form homodimer (EMD-44646; PDB 9BKV; **Fig. 3e**), a homodimeric complex comprising 1,614 residues. CryoZeta correctly placed atomic positions and achieved the highest sequence recall (0.92) among the compared methods; however, the overall TM-score was lower (0.631), indicating an incorrect global conformation. This discrepancy arises from a domain swap at residues 384–388 (boxed). The cryo-EM density in this region is locally low resolution, and consequently the reference PDB structure lacks coordinates for residues 379–383 in chain B. This example highlights that, despite locally accurate atomic placement, ambiguous density can lead CryoZeta to infer incorrect domain arrangements in homo-oligomeric targets.

### Models for Complexes with DNA and RNA

Next, we discuss four models of protein-nucleic acid complex targets (**Fig. 4**). The first example is the trimeric SenDRT9 RT-ncRNA complex (EMD-49525; PDB 9NLX; **Fig. 4a**), a homo-trimeric protein-RNA complex with 1,920 residues. CryoZeta accurately captured both the protein conformations and RNA positioning within the complex, achieving a TM-score of 0.983. In contrast, AF3 correctly modeled the individual protein but built less accurate RNA models and assembled them in incorrect relative orientations, resulting in a lower TM-score of 0.515. ModelAngelo correctly traced the protein trimer; however, the RNA was largely fragmented, yielding a TM-score of 0.669.

**Figure 4.**
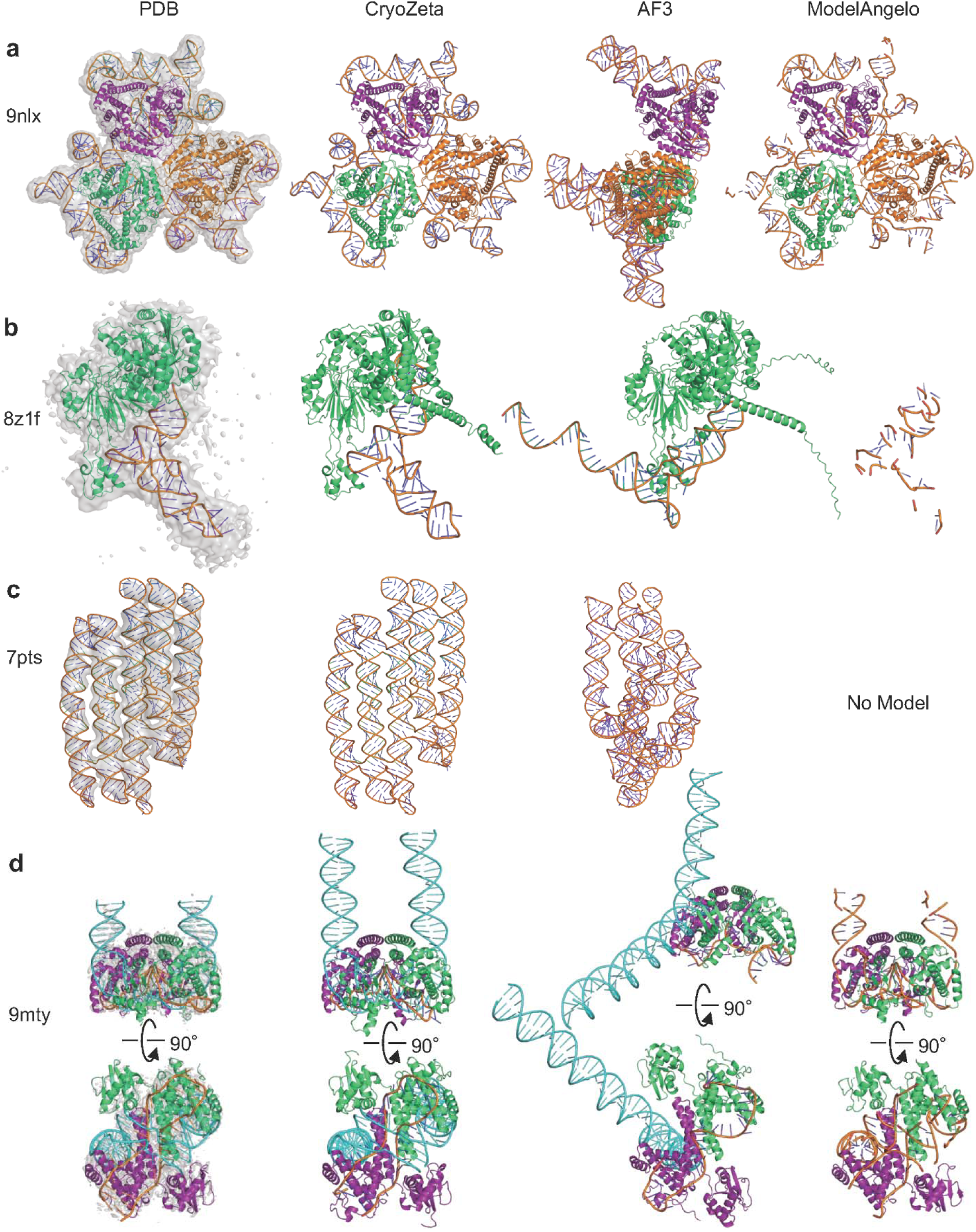
Examples of modeling for RNA, protein-RNA, protein-DNA-RNA complexes. For each target, the cryo-EM density map at the recommended level, the reference PDB structure, and models predicted by CryoZeta, AlphaFold3 (AF3), and ModelAngelo are shown. Modeling accuracy is evaluated using TM-score against the reference PDB structure. **a.** Trimeric SenDRT9 RT-ncRNA complex (EMD-49525, res. 3.20 Å; PDB 9NLX), a homo-trimeric protein–RNA complex with 1,920 residues. TM-scores are CryoZeta: 0.983; AF3: 0.515; and ModelAngelo: 0.669. **b.** Human ELAC2-tRNA complex (EMD-39726, res. 4.30 Å; PDB 8Z1F). This target is a monomeric protein–RNA complex with 912 residues. TM-scores are CryoZeta: 0.966; AF3: 0.883; and ModelAngelo: 0.0. **c**. RNA origami five-helix tile (EMD-13636, resolution 5.71 Å; PDB 7PTS), an RNA-only target consisting of 558 nucleotides. TM-scores are CryoZeta: 0.970; and AF3: 0.261. ModelAngelo failed to predict a model. **d.** IGR-TasR complex with tigRNA and target DNA after DNA cleavage (EMD-48616, res. 3.05 Å; PDB 9MTY), a homo-dimeric protein-DNA-RNA complex with 818 residues. DNA and RNA chains are colored by cyan and orange, respectively. TM-scores are CryoZeta: 0.989; AF3: 0.658; and ModelAngelo: 0.789.

The second example is the human ELAC2-tRNA complex (EMD-39726; PDB 8Z1F; **Fig. 4b**), a protein-RNA complex with 912 residues. CryoZeta generated almost perfect model with a TM-score of 0.966. AF3 produced an accurate protein structure model but made an incorrect RNA conformation (TM-score: 0.883), particularly at the anticodon position. For this low-resolution map (4.3 Å), ModelAngelo was unable to trace the protein and produced a fragmented RNA structure.

**Fig. 4c** is a 558 nucleotide-long single chain RNA target, an RNA origami five-helix tile (EMD-13636; PDB 7PTS). Despite the relatively low map resolution (res. 5.71 Å), CryoZeta accurately reconstructed the global conformation of the designed RNA structure, achieving a TM-score of 0.970. In contrast, AF3 produced a substantially less accurate RNA model (TM-score: 0.261), whereas ModelAngelo failed to generate a model for this target. This example highlights CryoZeta’s ability to directly model RNA-only structures from cryo-EM density maps, even for artificially designed RNA assemblies.

The last example is the IGR-TasR complex with tigRNA and target DNA after DNA cleavage (EMD-48616; PDB 9MTY; **Fig. 4d**), a homo-dimer protein complexed with DNA and RNA complex. CryoZeta produced the correct conformations of the protein, RNA, and DNA chains, including key DNA-RNA interfacial structure features associated with the DNA cleavage step mediated by the IGR-TasR system (TM-score: 0.989). The side and bottom views highlight that CryoZeta uniquely reconstructed the open DNA conformation and its interactions with RNA, capturing the key DNA cleavage process in this system. These DNA–RNA interactions were not reproduced by the other methods. In the AF3 model, individual protein structures were correct, but nucleic acid interactions were not captured correctly. AF3 also incorrectly modeled the DNA as a regular double-stranded helix. ModelAngelo traced the protein structure correctly but could not identify and generate the DNA structure (TM-score: 0.789).

### Modeling Large Complexes

Although CryoZeta was originally designed for targets up to 2,000 amino acids, we extended it with an iterative, chain-wise inference pipeline for larger complexes, integrating sequential structure prediction with adaptive density registration (Methods). To evaluate this protocol, we constructed a dataset of 51 targets with total sizes ranging from 2,000 to 5,000 and benchmarked it in comparison with AF3 (**Supplementary Table S14**). CryoZeta clearly outperformed AF3, achieving an average TM-score of 0.63 compared with 0.53 for AF3 (**Supplementary Fig. S12**). Four representative examples are shown in **Fig. 5**. Models by other methods are shown in **Supplementary Fig. S13.**

**Figure 5.**
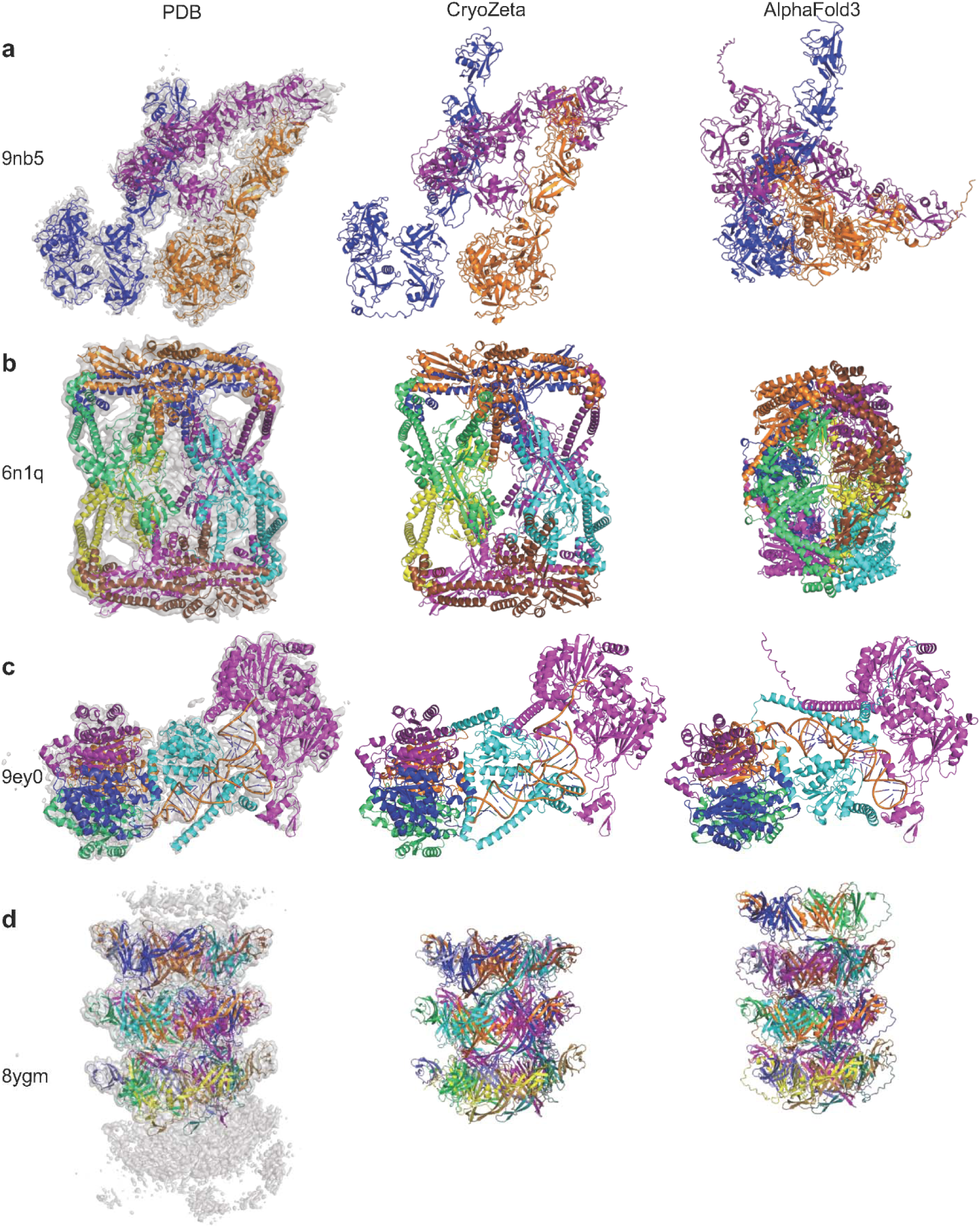
Examples of modeling for large protein and protein-RNA complexes. For each target, the cryo-EM density map at the recommended level, the reference PDB structure, and models predicted by CryoZeta, and AlphaFold3 (AF3) are shown. Modeling accuracy is evaluated using TM-score against the reference PDB structure. **a.** Autoinhibitory CD163 trimer (EMD-49213, res. 3.00 Å; PDB 9NB5), a homo-trimer complex with 3,036 residues. TM-scores are CryoZeta: 0.942 and AF3: 0.310. **b.** Dihedral oligomeric complex of the *Escherichia coli* GyrA N-terminal fragment (EMD-9317, res. 5.16 Å; PDB 6N1Q), a homo-octamer complex with 4,088 residues. TM-scores are CryoZeta: 0.982; and AF3: 0.448. **c.** Human mitochondrial RNase Z with tRNA-His (EMD-50050, res. 2.78 Å; PDB 9EY0), a hetero-hexameric protein–RNA complex with 2,268 residues. TM-scores are CryoZeta: 0.993; and AF3: 0.676. **d.** *Bacillus subtilis* A29 bacteriophage SPR tail tube protein (EMD-39256, res. 3.43 Å; PDB 8YGM), a homo-18 mer structure with 4,752 residues. TM-scores are CryoZeta: 0.977; and AF3: 0.657.

The first example is the autoinhibitory CD163 trimer (EMD-49213, res. 3.0 Å; PDB 9NB5; **Fig. 5a**), a homo-trimer protein complex with 3,036 residues. Each chain consists of eight domains. CryoZeta accurately reconstructed the overall architecture and inter-chain orientations of this large, multi-domain complex structure (TM-score: 0.942). In contrast, AF3 produced a more compact structure, which caused substantial errors in the relative orientations of the chains (TM-score: 0.310).

The next example is an even larger structure, a homo-octamer complex with 4,088 residues. (**Fig. 5b**; Dihedral oligomeric complex of GyrA N-terminal fragment; EMD-9317, res. 5.16 Å; PDB 6N1Q). Although the map is in a relatively low resolution, CryoZeta successfully reconstructed the overall architecture with high accuracy (TM-score: 0.982). AF3 again generated a compact structure, failing to reproduce the correct dihedral oligomeric complex (TM-score: 0.448).

**Fig. 5c** is the human mitochondrial RNase Z complexed with tRNA-His (EMD-50050, res. 2.78 Å; PDB 9EY0), a hetero-hexameric protein-RNA assembly with 2,268 residues. CryoZeta accurately reconstructed the protein-RNA interaction that is critical for recognition of the 5′ end of the mitochondrial tRNA (TM-score: 0.993). AF3 failed to model the RNA conformation correctly, producing incorrect protein–RNA interactions with less accurate protein–protein interfaces, resulting in a lower TM-score of 0.676.

The final example in **Fig. 5d** is a large homo 18-mer complex with 4,752 residues, *Bacillus subtilis* A29 bacteriophage SPR tail tube protein (EMD-39256, res. 3.43 Å; PDB 8YGM). CryoZeta accurately reconstructed the tail tube complex structure with a high TM-score of 0.977. AF3 also generated a tube-shape complex with substantially lower accuracy (TM-score = 0.657). In AF3 model, each layer consists of five chains rather than the six chains in the native structure resulting in a mediocre TM-score of 0.657. Among models by other methods (**Supplementary Fig. S13**), DiffModeler correctly recovered the native inter-chain contacts and overall architecture of the tail tube complex (TM-score: 0.961).

## Discussion

In this work, we developed CryoZeta, a unified structure modeling framework built upon the AF3 deep neural network architecture that incorporates cryo-EM density maps as an additional information source within a diffusion-based modeling paradigm. This design provides a principled integration of advances in cryo-EM–based structure modeling and macromolecular structure prediction. CryoZeta accurately models not only protein-only assemblies but also complexes containing RNA and DNA, and it consistently outperforms existing methods with substantial margins. Because CryoZeta inherits the AF3 architecture, it is also capable of modeling bound ligands, although a systematic evaluation of ligand modeling remains future work. Beyond single-state reconstruction, structural heterogeneity^6^ represents a major task in cryo-EM modeling. The integration of heterogeneous map reconstruction with atomic-level structure modeling is not well addressed here and remains an important direction for future methodological development.

## Acknowledgements

This work was partly supported by the National Institutes of Health (R35GM158267, R21AI187928) to DK and the National Science Foundation (IIS2211598, DMS2151678, DBI2146026 and DBI2422620 to DK and DBI2433490 to GT).

## Code Availability

CryoZeta is available as a web server at https://em.kiharalab.org/algorithm/CryoZeta.

## Author contributions

Z.Z., S.L. contributed to AF2-based code implementation. Z.Z., B.L., S.L., Y.Z. (in an early phase; Dec/2023–Apr/2024) contributed to AF3-based code implementation. Z.Z conducted model training for structure modeling. F.F. conducted model training and code implementation for U-net. F.F., S.L, N.I. contributed to U-Net optimization. S.L, Y.K., T.N, G.T contributed to dataset preparation. Z.Z, B.L, Y.K., G.T contributed to benchmarking against other methods. H.Z. assisted with optimization of structure fitting. M.K.K. assisted with code optimization. Z.Z., B.L., S.L., F.F., N.I, Y.K., G.T drafted the paper. G.T. prepared figures of model examples. Y.K. and B.L. implemented it in the EM web server. D.K. edited it.

## Conflict of Interests

GT and DK are founding members of Intellicule, LLC.

## Methods

### Dataset Construction

The datasets used in this study were first collected from PDB entries determined by cryo-EM with resolutions up to 8.0 Å, with release dates up to October 1, 2024. This initial query yielded 19,935 entries, which were then subjected to a series of rigorous filtering steps.

First, to ensure that the atomic structures in PDB entries and the corresponding maps match sufficiently well with each other, we computed cross-correlation (CC) coefficients, CC_box and CC_mask, between them using our in-house script. CC_box represents CC between the entire experimental map and a map simulated from the atomic structure at the resolution of the corresponding map, while CC_mask is CC computed only within the region occupied by the modeled macromolecules. Entries with either CC_box < 0.65 or CC_mask < 0.65 were discarded. We also excluded large macromolecules because they do not fit to GPUs of our computers. Entries with a bioassembly mmCIF file larger than 500 MB or a map file larger than 1 GB, as well as complexes of annotated with global symmetries of “Icosahedral”, “Tetrahedral”, or “Octahedral” were removed. Maps with non-orthogonal axes were also removed. Furthermore, entries containing unknown amino acid residues, structures with too large chain breaks, i.e. proteins with Cα-Cα distances ≥ 10 Å for consecutive amino acids, or nucleic acids with P-P distances ≥ 20 Å between consecutive nucleotides, were also filtered out. Finally, protein chains shorter than 30 residues and nucleic acid chains shorter than 50 residues were removed. After these filtering steps, 15,212 entries remained for clustering.

To ensure non-redundant dataset splits for training and testing, a two-stage clustering strategy was implemented. First, sequence clustering was performed on individual chains from PDB entries using MMseqs2^42^, with identity thresholds of 40 % for proteins and 80 % for nucleic acids. Then, two maps were grouped together if the maps contain at least a pair of sequences from the same cluster. This rigorous clustering approach resulted in a total of 1,383 unique map groups.

These map groups were then partitioned to create the final dataset splits. 1,106 clusters were assigned to the training set, 138 to the validation set, and 139 to the test set. This splitting methodology ensures that the resulting datasets are distinct not only at the individual sequence level but also at the level of whole macromolecular assemblies that might share homologous components.

To the datasets constructed initially, we added 4,809 maps on June 30, 2025. These maps met one of two criteria: either a resolution finer than 15 Å or a release date after October 1, 2024. These entries were processed using the identical filtering protocol as the original dataset. The filtered structures were assigned to the 1,383 existing map groups if they satisfy the similarity criteria mentioned above. The remaining unassigned entries were subsequently grouped, yielding 1184 entries in 484 new map groups. Due to large model complexity of CryoZeta, our model only support sequence length up to 2,000 on NVidia 80GB A100 GPU. This further reduced our validation set to 186 entries in 65 groups, and the test set to 441 entries in 200 groups.

The training sets for CryoZeta and CryoZeta-Interpolate were different, since large maps run into the memory issue when generating interpolation features between every pair of support points. To train CryoZeta, we use the full training set of 13,842 entries, which includes 10,881 protein complexes and 2,961 protein-nucleic acids complexes (**Supplementary Table S15**). To train CryoZeta-Interpolate, we used a training set of 9,685 entries, which includes 8,076 protein complexes and 1,609 protein-nucleic acids complexes (**Supplementary Table S16**).

### Training data preparation for the U-Net models for atom detection

For training the U-Net models, we used 6,103 maps in total, with 5,917 maps used for training and 186 maps for validation. These maps were a subset of from the 15,212 entries resulted by the filtering process mentioned above, selected based on the constraint of at most 15,000 amino acids/nucleotides. Specifically, the training dataset included 5,885 maps containing protein and 982 maps containing DNA/RNA (with 184 and 45 maps, respectively, for the validation split). To evaluate the performance of these models, we employed the same test dataset (200 maps) used for the CryoZeta main pipeline. There is no overlap between the test dataset and the training/validation datasets of the U-Net models, i.e. none of map pairs from training/validation sets and the test set belong to the same map group.

Maps in the datasets underwent preprocessing, which included resampling the map voxels to 1 Å with trilinear interpolation and normalization of their density values between 0 and 1. Then for each grid position of the maps, labels of atoms and residues, nucleotides were assigned based on the corresponding PDB files. For the atom detection network (**Supplementary Fig. S1**), we assigned labels for 8 atom types, 4 for proteins and 4 for nucleic acids. For proteins, we consider Cα atoms, backbone atoms (C, N, and O), Cβ atoms, and side-chain atoms. For nucleic acids, we put labels for C1′ atoms, sugar atoms (C2′, O2′ for RNA, C3′, O3′, C4′, O4′, C5′), phosphate atoms (P, OP1, OP2, O5′, OP3′), and base atoms. For the residue/nucleotide detection network, we assigned labels for 24 different classes: 20 amino acids and 4 nucleic-acid bases (with thymine and uracil merged into a single class). This assignment was done based on the closest atom or residue within 2 Å radius of the grid. Due to memory constraints while training, we cropped the map into boxes of size 48 x 48 x 48 grid points with a stride of 24×24×24. We discarded boxes in maps if the number of atom or residue labels was less than 0.1% of the total number of grid points in the box.

### Atom and residue/nucleotide detection by the U-Net module

Atoms and amino acid residues/nucleotides are detected by a network that connects two different U-Net models. The first one detects the atoms (8 classes, 4 for proteins and 4 for nucleic acids) and the second model detects the residues (24 classes, 20 for proteins and 4 for nucleic acids). Aside from the target labels, the training data for the two U-Net models were identical, i.e., both models were trained on the same maps, with only the output labels adjusted to atoms or residues, respectively. A higher-level architecture is provided in **Supplementary Fig. S1** and the individual U-Net architecture is shown in **Supplementary Fig. S2**. The network classifies each input voxel into specific categories, identifying 8 types of atoms or 24 types of residues within the input volume along with the background.

Both the atom and residue prediction models were trained for 30 epochs using a learning rate of 10^-3^ along with a cosine annealing scheduler. The loss function combined cross-entropy and dice loss, optimized with AdamW using default hyperparameters. Due to substantial class imbalance, class-specific weights were applied during loss computation. In the atom detection model, protein backbone and side-chain atoms were assigned a lower weight of 0.33, while Cα and Cβ atoms were given a higher weight of 2. For nucleic acids, C1′ atoms received a very high weight of 10 due to their rarity and importance for support point estimation, whereas base, phosphate, and sugar atoms had weights of 1, 1, and 0.5, respectively. For the residue detection model, weights of 1 and 2 were used for amino-acid residues and nucleic-acid bases.

### Assessment of voxel, atom, residue/nucleotide-level accuracy

Since detection is assigned to each voxel in a map, performance can first be evaluated at the voxel level and then propagated to the atom and residue/nucleotide levels according to an assignment rule. For each voxel, the atom closest to it within a 2 Å radius is assigned as the atom label for that voxel. The residue containing that atom is then assigned as the residue label for the voxel. For predictions, the atom or residue with the highest probability value for a voxel is considered the predicted label of that voxel. For an atom (e.g., Cα), voxel-level precision, recall, and F1-score are defined as follows. The definitions for voxel-level residue/nucleotide detection are analogous.

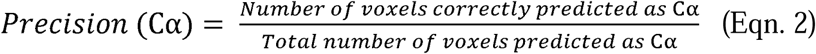

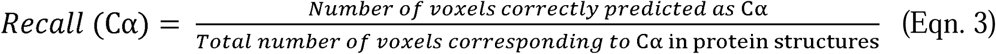

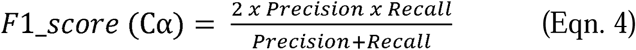

At the atom level, an atom is considered correctly predicted if more than 50% of its corresponding voxels are correctly predicted. Each voxel is assigned the label of the atom closest to it within a 2 Å radius. For prediction, a voxel is assigned to Cα or C1′ if their predicted probabilities exceed 0.25 and 0.2, respectively. These thresholds were determined based on the validation set. The definition of atom hit accuracy for Cα is shown below, and the same definition applies to RNA C1′ and DNA C1′.

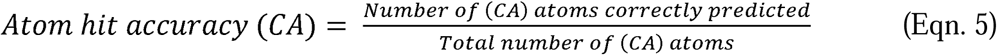

### Training the main network

The main network of CryoZeta (**Fig. 1**) was initialized with pre-trained Protenix weights^43^ except for networks related to cryo-EM, which was randomly initialized. Due to memory limitations, we used a cropped size of 512 residues. All loss terms used in AF3 were used. These include the MSE loss, distogram loss, predicted local distance difference (pLDDT) loss, pairwise atom-atom aligned error (PAE), predicted distance error (PDE), and experimentally resolved loss, where the network is trained to predict whether an atom is experimentally resolved. We adopted the weights of the loss terms based on the AF3 paper. In addition, we used point-residue distogram loss and point-noise loss with a weight of 1 to supervise cryo-EM information. Point-residue distogram loss helps the model to learn the relationships between predicted support points and Cα atoms (or C1′ atoms in the case of nucleotides). The point-residue distogram loss is defined as:

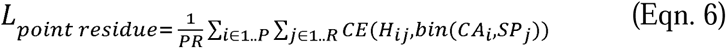

where P is the number of support points, R is number of residues/nucleotides in the native structure, H ∈ PxRx10 is the logits predicted by the model that is linearly projected from the point-residue representation, CA and SP is the Cα/C1′ atoms in native structure and support point, accordingly, and CE is the cross entropy loss. During training, the support point-residue representations were linearly projected into 10 distance bins, 0-0.5 Å, 0.5-1 Å, 1-2 Å, 2-3 Å, 3-5 Å, 5-7.5 Å, 7.5-10 Å, 10-12.5 Å, 12.5-15 Å, and >15 Å. The ground truth labels for point-residue loss was calculated by computing the Euclidean distance between Cα/C1′ in the native structure and detected support points.

The point noise loss was used to help the model separate noisy supports point from other support points. Here we define a noisy support point as a support point that is at least 5 Å away to any Cα/C1′ atom in the native structure. The point-noise loss is defined as:

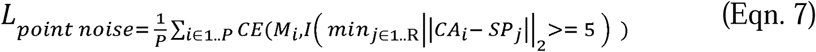

where P is number of support points, R is number of residues/nucleotides in the native structure, M ∈ P is the logits predicted by the model that is linearly projected from the point-residue representation, CA and SP is the Cα/C1′ atoms in the native structure and support point, accordingly, and CE is the cross entropy loss. All other training related parameters, such as learning rate and diffusion sample steps, were inherited from Protenix.

### Structure fitting to a map

To align predicted structures to cryo-EM maps, we implemented two complementary point-cloud registration strategies: (i) an SVD-based Kabsch alignment^44^ using predicted point–residue correspondences, and (ii) TEASER++^45^ followed by Generalized Iterative Closest Point (GICP^46,47^) refinement. For the first strategy, given a predicted point–residue distance distribution and confidence scores, we first identify all point–residue pairs predicted to be within 5Å (Cα and C1′ are used as main atoms to represent the residues and nucleotides). For each candidate structure, we superimpose the residue coordinates onto their corresponding support point coordinates using the SVD-based Kabsch alignment algorithm. We perform the algorithm twice using two point–residue distance confidence thresholds, 0.8 and 0.4. This is because sometimes the higher confidence threshold selects too few point–residue pairs for stable fitting. The lower confidence threshold selects more pairs with some accuracy trade-off. The resulting rotation and translation are then applied uniformly to all atoms of the structure. However, it is possible that SVD does not provide the optimal superimposition. This is because sometimes the point-distogram prediction may not be confident enough for large maps, resulting in poor selection of point-residue pair for super imposition. In such case, we adopt two-stage alignment strategy where we first use a point-cloud matching method TEASER++ to fit the structure and we use GICP to refine the fitted result. Different from SVD where support point needs to be filtered to match the number of CA/C1′ atoms, TEASER++ directly takes the whole support points as input and computes the optimal rotation and translation. After the first rough initial superimposition by TEASER++, we then use GICP to align the structure more closely. Such two-stage alignment strategy has also been used in aligning cryo-EM maps and structures^48^. The quality of the superimposed structure is evaluated by a recall metric defined as the fraction of main atoms that lie within 3Å of any support point, and the transformation yielding the highest recall is selected as the final fit. In rare cases where both the SVD and TEASER++ algorithms yield zero recall, we apply VESPER^49^ fitting as a final measure.

### Modeling large complexes

To construct structures of large macromolecular complexes, we developed an iterative, chain-wise inference pipeline that combines sequential prediction with adaptive density registration. In this protocol, a multi-chain complex is decomposed into individual chains ordered by decreasing sequence length, and one copy of a chain is predicted per iteration, enabling progressive assembly. At each iteration, the predicted chain is registered to the cryo-EM density using the fitting strategy described above. Following each successful registration, we update the density representation by masking all support points within a 5.0 Å radius of the registered Cα/C1′ coordinates. This step removes resolved density and forces subsequent iterations to focus on unresolved regions. This iterative registration–filtering cycle reduces inter-chain interference and improves reconstruction fidelity for large macromolecular assemblies.

### Running existing methods for comparison

For AlphaFold3, the publicly available AlphaFold3 codebase (v3.0.1) available at https://github.com/google-deepmind/alphafold3 was used, following the standard inference protocols recommended by the developers. Predictions were generated using the full-length sequences of the target complexes without providing any templates or custom MSAs beyond those automatically generated by the standard AlphaFold3 pipeline. Five structures were generated using a single randomly chosen seed for each target.

For ModelAngelo, we used the official ModelAngelo codebase release v1.0.14 available at https://github.com/3dem/model-angelo. The input for ModelAngelo consisted of the experimental cryo-EM maps along with the corresponding protein and nucleic acid sequences for each chain present in the target complex. Default parameters for ModelAngelo were used throughout the model building process.

For DiffModeler, the public codebase release v2.3 available at https://github.com/kiharalab/DiffModeler was used. The initial structural models of proteins were predictions generated by AlphaFold3 for the corresponding target complexes. We used a half of author recommended contour level as input parameter.

For CryFold, we used the publicly released v1.3 version available at https://github.com/SBQ-1999/CryFold. CryFold takes a cryo-EM density map together with its corresponding protein sequence in the FASTA format as input. Accordingly, we obtained all maps from the EMDB and downloaded matching sequences from the PDB. We ran all predictions using the default inference settings provided in the documentation.

For Protenix, we benchmarked the official v0.0.1 release available at https://github.com/bytedance/Protenix. Input modalities included protein, DNA, RNA, and small-molecule ligands, all parsed directly from mmCIF files downloaded from the PDB. We deliberately omitted optional modified residue annotations and covalent bond definitions. This Protenix version operates in a template-free mode and does not support RNA/DNA MSAs, thus we supplied only protein MSAs. Inference was run with the default hyperparameters with four Pairformer recycles and 20 diffusion timesteps.

We used DeepMainmast v1.0.0 obtained at https://github.com/kiharalab/DeepMainMast. We performed the full pipeline of DeepMainmast, including full-atom modeling using Rosetta. Predicted models by AlphaFold3 were used as external structural information. The contour level of the cryo-EM map was set to half of the value recommended by the authors.

## Notes

### Competing Interest Statement

GT and DK are founding member of Intellicule LLC.

## References

1 Yip, K. M., Fischer, N., Paknia, E., Chari, A. & Stark, H. Atomic-resolution protein structure determination by cryo-EM. Nature 587, 157–161 (2020). 10.1038/s41586-020-2833-4

2 Murata, K. & Wolf, M. Cryo-electron microscopy for structural analysis of dynamic biological macromolecules. Biochim Biophys Acta Gen Subj 1862, 324–334 (2018). 10.1016/j.bbagen.2017.07.020

3 Casanal, A., Lohkamp, B. & Emsley, P. Current developments in Coot for macromolecular model building of Electron Cryo-microscopy and Crystallographic Data. Protein Sci 29, 1069–1078 (2020). 10.1002/pro.3791

4 Adams, P. D. et al. PHENIX: a comprehensive Python-based system for macromolecular structure solution. Acta crystallographica. Section D, Biological crystallography 66, 213–221 (2010). 10.1107/S0907444909052925

5 Croll, T. I. ISOLDE: a physically realistic environment for model building into low-resolution electron-density maps. Acta Crystallogr D Struct Biol 74, 519–530 (2018). 10.1107/S2059798318002425

6 Farheen, F., Terashi, G., Zhu, H. & Kihara, D. AI-based methods for biomolecular structure modeling for Cryo-EM. Current opinion in structural biology 90, 102989 (2025). 10.1016/j.sbi.2025.102989

7 Li, S., Terashi, G., Zhang, Z. & Kihara, D. Advancing structure modeling from cryo-EM maps with deep learning. Biochem Soc Trans 53, 259–265 (2025). 10.1042/BST20240784

8 Zhu, H., Terashi, G., Farheen, F., Nakamura, T. & Kihara, D. AI-based quality assessment methods for protein structure models from cryo-EM. Curr Res Struct Biol 9, 100164 (2025). 10.1016/j.crstbi.2025.100164

9 Terashi, G. & Kihara, D. De novo main-chain modeling for EM maps using MAINMAST. Nature communications 9, 1618 (2018). 10.1038/s41467-018-04053-7

10 Chen, M., Baldwin, P. R., Ludtke, S. J. & Baker, M. L. De Novo modeling in cryo-EM density maps with Pathwalking. Journal of structural biology 196, 289–298 (2016). 10.1016/j.jsb.2016.06.004

11 Pfab, J., Phan, N. M. & Si, D. DeepTracer for fast de novo cryo-EM protein structure modeling and special studies on CoV-related complexes. Proceedings of the National Academy of Sciences of the United States of America 118 (2021). 10.1073/pnas.2017525118

12 Terashi, G., Wang, X., Prasad, D., Nakamura, T. & Kihara, D. DeepMainmast: integrated protocol of protein structure modeling for cryo-EM with deep learning and structure prediction. Nature methods 21, 122–131 (2024). 10.1038/s41592-023-02099-0

13 Zhang, X., Zhang, B., Freddolino, P. L. & Zhang, Y. CR-I-TASSER: assemble protein structures from cryo-EM density maps using deep convolutional neural networks. Nature methods 19, 195–204 (2022). 10.1038/s41592-021-01389-9

14 Chen, S. et al. Protein complex structure modeling by cross-modal alignment between cryo-EM maps and protein sequences. Nat Commun 15, 8808 (2024). 10.1038/s41467-024-53116-5

15 Wang, X., Terashi, G. & Kihara, D. CryoREAD: de novo structure modeling for nucleic acids in cryo-EM maps using deep learning. Nature methods 20, 1739–1747 (2023). 10.1038/s41592-023-02032-5

16 Jamali, K. et al. Automated model building and protein identification in cryo-EM maps. Nature 628, 450–457 (2024). 10.1038/s41586-024-07215-4

17 Wang, J. et al. End-to-end Cryo-EM complex structure determination with high accuracy and ultra-fast speed. Nature Machine Intelligence, 1–13 (2025).

18 Su, B., Huang, K., Peng, Z., Amunts, A. & Yang, J. Improved automated model building for cryo-EM maps using CryFold. in. bioRxiv 2024, 13.623164 (2024).

19 Li, P. et al. An end-to-end approach for protein folding by integrating Cryo-EM maps and sequence evolution. bioRxiv, 2023.2011. 2002.565403 (2023).

20 Jumper, J. et al. Highly accurate protein structure prediction with AlphaFold. Nature 596, 583–589 (2021). 10.1038/s41586-021-03819-2

21 Zhang, Z. et al. DEMO-EM2: assembling protein complex structures from cryo-EM maps through intertwined chain and domain fitting. Brief Bioinform 25 (2024). 10.1093/bib/bbae113

22 Frigo, M. & Johnson, S. G. The design and implementation of FFTW3. Proceedings of the IEEE 93, 216–231 (2005).

23 He, J., Lin, P., Chen, J., Cao, H. & Huang, S. Y. Model building of protein complexes from intermediate-resolution cryo-EM maps with deep learning-guided automatic assembly. Nat Commun 13, 4066 (2022). 10.1038/s41467-022-31748-9

24 Wang, X., Zhu, H., Terashi, G., Taluja, M. & Kihara, D. DiffModeler: large macromolecular structure modeling for cryo-EM maps using a diffusion model. Nat Methods 21, 2307–2317 (2024). 10.1038/s41592-024-02479-0

25 Ho, J., Jain, A. & Abbeel, P. Denoising diffusion probabilistic models. Advances in neural information processing systems 33, 6840–6851 (2020).

26 Abramson, J. et al. Accurate structure prediction of biomolecular interactions with AlphaFold 3. Nature 630, 493–500 (2024). 10.1038/s41586-024-07487-w

27 Raghu, R., Levy, A., Wetzstein, G. & Zhong, E. D. Multiscale guidance of AlphaFold3 with heterogeneous cryo-EM data. arXiv preprint arXiv:2506.04490 (2025).

28 Zhang, Y. & Skolnick, J. Scoring function for automated assessment of protein structure template quality. Proteins 57, 702–710 (2004). 10.1002/prot.20264

29 Mirdita, M. et al. ColabFold: making protein folding accessible to all. Nat Methods 19, 679–682 (2022). 10.1038/s41592-022-01488-1

30 UniProt, C. UniProt: the Universal Protein Knowledgebase in 2025. Nucleic Acids Res 53, D609–D617 (2025). 10.1093/nar/gkae1010

31 Zhang, C., Zhang, Y. & Pyle, A. M. rMSA: A Sequence Search and Alignment Algorithm to Improve RNA Structure Modeling. J Mol Biol 435, 167904 (2023). 10.1016/j.jmb.2022.167904

32 Consortium, R. N. RNAcentral in 2026: genes and literature integration. Nucleic Acids Res (2025). 10.1093/nar/gkaf1329

33 Ontiveros-Palacios, N. et al. Rfam 15: RNA families database in 2025. Nucleic Acids Res 53, D258–D267 (2025). 10.1093/nar/gkae1023

34 Sayers, E. W. et al. Database resources of the national center for biotechnology information. Nucleic Acids Res 50, D20–D26 (2022). 10.1093/nar/gkab1112

35 Huang, H. et al. in ICASSP 2020-2020 IEEE International Conference on Acoustics, Speech and Signal Processing (ICASSP). 1055–1059 (IEEE).

36 Carreira-Perpinan, M. A. in 2006 IEEE Computer Society Conference on Computer Vision and Pattern Recognition (CVPR’06). 1160–1167 (IEEE).

37 Bekker, G. J. et al. Protein Data Bank Japan: Computational Resources for Analysis of Protein Structures. J Mol Biol 437, 169013 (2025). 10.1016/j.jmb.2025.169013

38 ww, P. D. B. C. EMDB-the Electron Microscopy Data Bank. Nucleic acids research 52, D456–D465 (2024). 10.1093/nar/gkad1019

39 Zhang, C., Shine, M., Pyle, A. M. & Zhang, Y. US-align: universal structure alignments of proteins, nucleic acids, and macromolecular complexes. Nat Methods 19, 1109–1115 (2022). 10.1038/s41592-022-01585-1

40 Liebschner, D. et al. Macromolecular structure determination using X-rays, neutrons and electrons: recent developments in Phenix. Acta Crystallogr D Struct Biol 75, 861–877 (2019). 10.1107/S2059798319011471

41 Terashi, G., Wang, X., Maddhuri Venkata Subramaniya, S. R., Tesmer, J. J. G. & Kihara, D. Residue-wise local quality estimation for protein models from cryo-EM maps. Nature Methods 19, 1116–1125 (2022). 10.1038/s41592-022-01574-4

42 Steinegger, M. & Soding, J. MMseqs2 enables sensitive protein sequence searching for the analysis of massive data sets. Nature biotechnology 35, 1026–1028 (2017). 10.1038/nbt.3988

43 Team, B. A. A. S. et al. Protenix-advancing structure prediction through a comprehensive AlphaFold3 reproduction. BioRxiv, 2025.2001. 2008.631967 (2025).

44 Kabsch, W. A solution for the best rotation to relate two sets of vectors. Foundations of Crystallography 32, 922–923 (1976).

45 Yang, H., Shi, J. & Carlone, L. Teaser: Fast and certifiable point cloud registration. IEEE Transactions on Robotics 37, 314–333 (2020).

46 Segal, A., Haehnel, D. & Thrun, S. in Robotics: science and systems. 435 (Seattle, WA).

47 Koide, K. small_gicp: Efficient and parallel algorithms for point cloud registration. Journal of Open Source Software 9, 6948 (2024).

48 He, B. et al. Accurate global and local 3D alignment of cryo-EM density maps using local spatial structural features. Nat Commun 15, 1593 (2024). 10.1038/s41467-024-45861-4

49 Han, X., Terashi, G., Christoffer, C., Chen, S. & Kihara, D. VESPER: global and local cryo-EM map alignment using local density vectors. Nature communications 12, 2090 (2021). 10.1038/s41467-021-22401-y

